# Transferability of ecological forecasting models to novel biotic conditions in a long-term experimental study

**DOI:** 10.1101/2023.11.01.565145

**Authors:** Patricia Kaye T. Dumandan, Juniper L. Simonis, Glenda M. Yenni, S. K. Morgan Ernest, Ethan P. White

**Author notes:** **Corresponding author**: S. K. Morgan Ernest.

## Abstract

Ecological forecasting models play an increasingly important role for managing natural resources and assessing our fundamental knowledge of processes driving ecological dynamics. As global environmental change pushes ecosystems beyond their historical conditions, the utility of these models may depend on their transferability to novel conditions. Because species interactions can alter resource use, timing of reproduction, and other aspects of a species’ realized niche, changes in biotic conditions, which can arise from community reorganization events in response to environmental change, have the potential to impact model transferability. Using a long-term experiment on desert rodents, we assessed model transferability under novel biotic conditions to better understand the limitations of ecological forecasting. We show that ecological forecasts can be less accurate when the models generating them are transferred to novel biotic conditions, and that the extent of model transferability can depend on the species being forecast. We also demonstrate the importance of incorporating uncertainty in forecast evaluation with transferred models generating less accurate and more uncertain forecasts. These results suggest that how a species perceives its competitive landscape can influence model transferability, and that when uncertainties are properly accounted for, transferred models may still be appropriate for decision making. Assessing the extent of the transferability of forecasting models is a crucial step to increase our understanding of the limitations of ecological forecasts.

**Open Research Statement:** Data (10.5281/zenodo.8436468) and code (10.5281/zenodo.11053744) used to conduct data analyses are archived in Zenodo and supplementary results (10.6084/m9.figshare.25733691.v1) in Figshare.

## INTRODUCTION

Ecological forecasts - predictions for the future state of ecosystems - are increasingly important for understanding, managing, and conserving natural and managed systems (Dietze et al., 2018; Bodner et al., 2021; Lewis et al., 2023). Most ecological forecasts are based on models that are fit to the dynamics of the ecosystem being studied. Making forecasts from these models assumes that the behavior of the ecosystem will remain the same in the future. However, with climate change, land use change, and the spread of invasive species, many ecosystems will be experiencing environmental shifts making it unclear how forecasting models will perform as altered conditions take effect (Yates et al. 2018). Deploying models in novel conditions is also important in ecology because data limitations often require us to use information from one ecosystem to make inferences and predictions for less well-studied ecosystems (e.g., Fitzpatrick and Hargrove 2009, McCune 2016). Evaluating forecasting models in novel conditions can also provide an assessment of the generality of ecological theory (Lewis et al. 2023), strengthening our overall knowledge of how ecological systems operate. Therefore, a crucial step for ecological forecasting, and ecology more broadly, is understanding how well models perform under conditions that differ from those used for model development (Werkowska et al. 2017; Yates et al., 2018; Charney et al. 2021; Lewis et al., 2023).

The effectiveness of models for making predictions under novel conditions is known as model transferability (Randin et al. 2006). In ecology, novel conditions in space or time can result from differences in abiotic conditions, the biotic context (e.g., the presence or abundance of native or non-native species), or both (Radeloff et al 2015) While more studies of model transfer are necessary in ecological forecasting (Lewis et al. 2023), initial analyses indicate that model transferability is negatively influenced by model complexity (with more complex models tending to generalize less successfully than simpler models; Wenger and Olden 2012, Liu et al. 2020, Chen et al. 2023), and the degree of ecological novelty (with larger differences in environmental conditions resulting in poorer transfer; Sequeira et al 2018, Lewis et al. 2023). While analyses related to ecological novelty often focus on abiotic conditions or coarse biotic conditions such as habitat structure (e.g., Spence and Tingley 2020, Quiao et al 2019, Regos et al 2019), altered biotic conditions are also a potential concern for model transferability. Changes in the biotic conditions can fundamentally alter the observed dynamics between a species and their resources and environment (Casini et al. 2009, Tingley et al. 2014). For example, the loss of a key species may impact the surrounding habitat and therefore the abundance of other species (Power et al. 1996, Goheen et al. 2018), the loss of predators or competitors may relieve biotic pressures on species allowing them to increase in abundance (e.g., Holt et al. 2008, Trewby et al 2007, Leal et al 1998), and the arrival of invasive species may dramatically depress abundances through predation and competition that the resident species are not adapted to deal with (Wiles et al. 2003, Gallardo et al 2016). Shifts in the strength and number of species interactions can also impact the skill of forecasts (Daugaard et al. 2022). Thus, changes in biotic conditions can potentially alter the transferability of forecasting models by impacting the relationship between a species’ population dynamics and its environment, even if other environmental conditions remain unchanged. Because many environmental issues - i.e., climate-induced range shifts (Williams and Jackson 2007), colonization of invasive species (David et al. 2017), and global extinctions (Raffaelli 2004) - involve altered biotic conditions, understanding the impacts of biotic changes on forecasts is critical for understanding the potential limitations of model transferability for ecological forecasting.

Little is known about the impact of altered biotic conditions on model transferability in forecasting because suitable data is limited (Paniw et al. 2023). Community change - caused by extinction, colonization, or shifts in dominance - generally occurs with larger-scale changes in abiotic environment, habitat structure, or other landscape-level alterations (Whittaker 1960, Wiens and Rotenberry 1981, Gentry 1988). Thus, disentangling the effects of community change on model transferability from other environmental changes requires experimental manipulations that selectively manipulate species to generate different biotic communities experiencing the same general environment. Most experiments are short-term, however, lasting on average one to three years (Field et al. 2007, Magnusson 1990), which reduces the data available to both fit a model and test the model outcomes, especially if assessing performance under natural environmental variation is a goal. Therefore, to assess the impact of biotic shifts on model transferability and forecast performance, long-term experimental manipulations are required.

Here, we assess model transferability under novel biotic conditions using a long-term experiment on desert rodents in the southwestern US. For over 40 years, the Portal Project has collected monthly data on natural and experimentally manipulated rodent communities all experiencing the same abiotic environment. In this experiment a competitively dominant genus, *Dipodomys* spp. (kangaroo rats), has been excluded resulting in significant impacts on other species in the system (Brown 1998, Bledsoe and Ernest 2019, Diaz and Ernest 2022). Using this unique dataset, we investigate how biotic context influences parameter estimation and prediction accuracy when models are fit under one set of biotic conditions but used to forecast under a novel biotic regime. Changes in model parameterizations in response to biotic conditions would indicate that inferences about population dynamics under climate change are sensitive to the other species present in the ecosystem, while changes in forecast accuracy would indicate potentially important challenges for ecological forecasting.

## METHODS

### Rodent data

To examine whether shifting biotic conditions can impact model transferability we obtained data on rodent population dynamics from a long-term monitoring program in the Chihuahuan Desert near Portal, Arizona (Brown 1998, Ernest et al. 2018). The 20-ha study site consists of 24 50 m x 50 m plots, each enclosed with a 50 cm fence with different sized gates to manipulate rodent species access. Plots are randomly assigned to three levels of rodent community manipulation: controls (large gates, all rodents have full access to plots), kangaroo rat removals (small gates, behaviorally dominant seedeaters, *Dipodomys* spp., are excluded), and total rodent removals (no gates, all rodents excluded but occasional transient individuals occur). The rodent communities in each plot are censused monthly around the new moon using 49 Sherman traps, and basic information is collected for all trapped rodents. Further details about the experimental setup and sampling methods are discussed elsewhere (Ernest et al. 2016, Ernest et al. 2018). In this study, we only used data on the communities found in long-term (i.e., treatments maintained across all years) controls (plots 4, 11, 14, 17) and kangaroo rat removal (plots 3, 15, 19, 21). Data were obtained using the ‘portalr’ package (Christensen et al. 2019) and are also archived on Zenodo (10.5281/zenodo.8436468).

We used count data from long-term control and *Dipodomys* removal plots for the desert pocket mouse (*C. penicillatus*) and Bailey’s pocket mouse (*C. baileyi*). We selected these species because there were extended time periods when they were relatively abundant in both control and kangaroo rat removal plots (i.e., fewer zeros which can complicate modeling) and both species respond strongly to the experimental removal of *Dipodomys* (Bledsoe and Ernest 2019, Diaz and Ernest 2022). Previous modeling efforts (Christensen et al. 2018) found five different community regimes at the site, so we selected the two regimes where each non-*Dipodomys* species was highly abundant (Fig. 1). Regime transitions are probabilistic, so we used the edge of the range for the transition to ensure that the data was entirely within the regime and did not include transitions between the regimes. Continued trapping at the site suggests that the 2010-2015 regime has continued and so we extended this time period to the end of 2019, shortly before an extensive gap in data collection due to the COVID-19 pandemic. This resulted in data for *C. baileyi* spanning from December 1999 to June 2009 (new moon number 278-396) and for *C. penicillatus* from September 2010 to December 2019 (new moon number 411-526).

**Figure 1.**
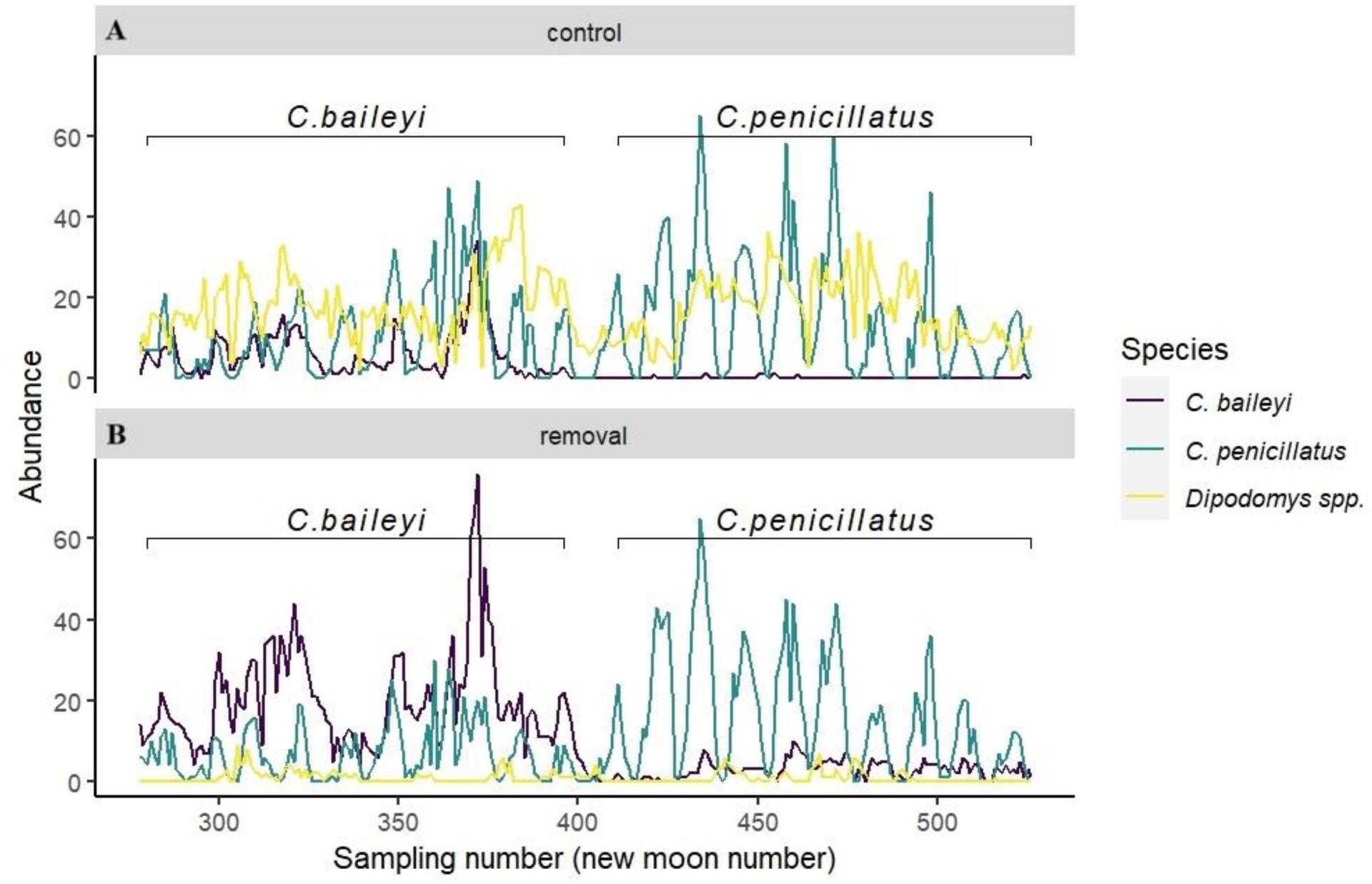
Time-series data on *Dipodomys* spp. (yellow lines), *Chaetodipus baileyi* (purple lines), and *C. penicillatus* (green lines) on control plots (A) and removal plots (B) in a long-term experiment near Portal, AZ. The two species-specific periods used for modeling and forecasting *Chaetodipus* spp. are indicated by brackets.

### Environmental covariates data

We used environmental covariates known to be important drivers of ecological processes in this desert ecosystem. We obtained site-level monthly data on mean air temperature (°C) and cumulative precipitation (millimeters) that fell during warm or cool months (calculated as the sum of precipitation that fell on days when minimum temperature was > or < 4 °C) through the ‘portalr’ package (Christensen et al. 2019). This data is collected by an on-site weather station and any gaps are filled with modeled data from nearby regional weather stations (Ernest et al. 2018). Mean air temperature is a strong driver of seasonal abundance of *C. penicillatus*, and potentially other smaller rodent species, as it influences foraging effort and seasonal activity (i.e., entering bouts of torpor or seasonal dormancy; Reynolds and Haskell, 1949, Meyer and Valone 1999). Lags are known to be important in the system, with one-month lags frequently capturing the time it takes for individuals to behaviorally respond to changing environmental conditions (Thibault et al. 2010,Clark et al., 2023)), so we used a one-month lag for the temperature response. We used cumulative precipitation over the preceding 365-day as a covariate because the size of granivore populations responds to precipitation-related changes in annual seed production over the last year, with little carryover to subsequent years (Brown et al. 1979, Brown and Heske 1990). In this ecosystem, winter and summer precipitation have different influences on plant growth and seed production, with cool precipitation being important for the winter annual plant community and shrub growth and establishment, and warm precipitation being important for the summer annual and perennial plant community (for information on the two mostly distinct annual plant communities at the site, see Ernest et al. 2018).

### Modeling Approach

To assess how well forecasting models can transfer to different biotic conditions, we fit models separately to the control plots (where kangaroo rats are present) and the *Dipodomys* removal plots (where kangaroo rats are absent). We fit these treatment-specific models for each species to allow us to compare the parameters of the models from the different treatments and assess how well the models from one treatment could predict abundances on the other treatment.

The general model structure was an autoregressive model with 1 time-step and 1-year lags plus the three environmental covariates. Each time-series model had the form:

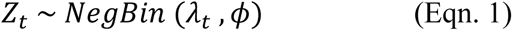

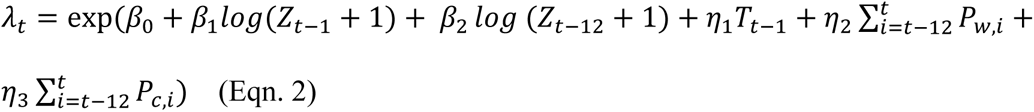

where *Z*_*t*_, the species-specific count at time *t* is drawn from a negative binomial distribution with parameters *λ*_*t*_(the conditional mean of abundance at time *t*) and *ϕ* (dispersion parameter). The negative binomial distribution was chosen to model discrete abundances while addressing overdispersion and provides a better fit to this dataset than the Poisson (Appendix S1 Table 1). The conditional mean was modeled as a function of an intercept (*β*_0_), autoregressive terms for the abundance of the previous observation (*β*_1_ *log*(*Z*_*t*−1_ + 1)) and the abundance at the same time in the previous year *β*_2_ *log*(*Z*_*t*−12_ + 1), i.e., 12 time steps), linear terms for the effects of mean temperature of the previous month (*η*_1_*T*_*t*−1_) and the annual cumulative values of warm 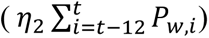 and cold 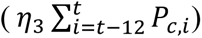 precipitation. One time-step and annual lags were selected to capture short-term correlations and seasonal cycles. These choices are supported by analyses of autocorrelation structure on both the empirical data and model residuals (Appendix S1 Figure 6). The annual lag is less important for *C. baileyi* but was kept to allow direct comparisons between species. The inclusion of weather data up to time *t* is realistic in the forecasting context of this system since the weather data is collected in real-time and automatically integrated into the dataset (White et al. 2019) making it available for predictions for time *t*. Models were implemented in a frequentist framework using the ‘tscount’ package (Liboschik et al. 2017) in R version 4.1.1 (R Core Team, 2021). We chose to only include time-series terms based on the observed counts (using *Z*_*t*−1_ and *Z*_*t*−12_ and not using *λ*_*t*−1_ and *λ*_*t*−12_) to allow models to be effectively transferred. This means that “internal” and “external” forms of the model (see Liboschik et al. 2017) are the same.

This modeling approach requires consistently sampled data (Liboschik et al. 2017), so values for the small number of missing samples (n = 9/116 (7.8%) for *C. penicillatus* and n = 4/119 (3.4%) for *C. baileyi*) were imputed using linear interpolation. Because we trap as close as possible to the new moon (Ernest et al. 2018) the annual periodicity of sampling is not exactly 12 periods. There are on average 12.37 new moons/year. Therefore, we also fit the models using a 13-period lag for comparison. We also evaluated models with both scaled and unscaled covariates. The results were qualitatively similar in both cases (Appendix S2 Figs 2-7). Code used to conduct analyses is archived on Zenodo (10.5281/zenodo.11053744).

To examine the relative importance of biotic conditions in driving variation in model parameters across the time-series, sequential model fitting with rolling origins was performed to generate a number of different forecasting models each with five years of training data (Simonis et al. 2021). Models were fit separately for data on *C. penicillatus* and *C. baileyi* in control and removal plots. We used the ‘rsample’ package (Frick et al. 2022) to conduct rolling origin modeling on each dataset, with 60 data points (12 observations/year for 5 years) used for model training and 12 data points (12 observations/year for 1 year) for model evaluation. This produced 45 sets of overlapping models and evaluations for *C. penicillatus* and 48 sets for *C. baileyi*.

### Comparing model parameters

We compared the distribution of the coefficients estimated from each rolling origin model to determine the degree of similarity in the coefficients between the treatment-specific datasets. Differences in coefficients are a prerequisite for differences in forecast performance between the two treatments when models are transferred, and quantifying any differences will provide insight into the aspects of biology that are impacting model transfer. We compared the estimated coefficients by quantifying the degree of overlap in the kernel density estimates of each parameter using the ‘overlapping’ package (Pastore et al. 2022). The resulting overlap index is on a scale from 0 to 1, with 0 indicating distinct distributions of estimated coefficients indicating a strong change and 1 indicating completely overlapping distributions indicating no change (Pastore et al. 2022). This analysis combines variation between the original non-transferred and transferred models for a single origin with variation within models among origins, providing perspective on whether the influence of biotic conditions is sufficiently strong to be observable even when temporal variation in parameter estimates is present. To also focus directly on the shift in estimated coefficients in response to the experimental manipulation of biotic context, we characterized the proportion of pairwise changes for each origin by calculating the difference in estimated coefficients from each treatment.

We checked to make sure that the interpretability of the parameters associated with individual environmental covariates was not unduly influenced by collinearity by performing pairwise correlation and covariance assessments among the covariates and their parameters. Environmental covariates used in the models had low correlations and the covariances and correlation values of their coefficients were low (Appendix S1).

### Model transfer

To assess model transferability to different biotic contexts, we generated forecasts for both the treatment data to which the model was fit (non-transferred model) and to the data for the other treatment (transferred model). Forecasts from transferred models (e.g., model parameters for the removal model used to predict counts in the control plots) were made using the initial conditions from time-series being forecast, and the model parameters for the data the model was trained on. Similar steps were followed to generate forecasts for the non-transferred model (where data and model were matched, e.g., control model used to predict control data).

### Forecasting evaluation

We evaluated the models from each rolling origin using end-sample evaluation - where forecasts are made for observations *h* time-steps past the end of the final observation in the training time-series and evaluating on the observed test data (Simonis et al. 2021). We made forecasts for three-time horizons (h = 1, approximately 1 month; h = 6, approximately half a year; and h = 12, approximately 1 year) for each rolling origin. The test data for each model were the subsequent 12 observations following each set of training data (following White et al. 2019). We assessed accuracy of point forecasts using root mean squared error (RMSE) and forecast uncertainty using Brier score, which is a proper scoring rule that extends the mean squared error to distributional forecasts (Simonis et al. 2021). For each species, RMSE and Brier scores of non-transferred and transferred models were calculated for each rolling origin model at each forecast horizon. We then calculated the difference between the pairs of RMSE values and Brier scores from each origin for the non-transferred and transferred models to assess the effect of novel biotic conditions in driving forecast predictability. Negative values for RMSE and Brier score differences indicate better forecast performance from the non-transferred model, and positive values indicate better forecast performance from the transferred model.

## RESULTS

### Model parameter comparison

The two species differed in whether their model parameters were influenced by the biotic context. For *C. penicillatus*, the estimated coefficients did not differ significantly for models fit to data on control and removal plots, indicating similar associations between abundances and environmental variables in both plot types (Fig. 2). This is indicated by the relatively high overlap in the distributions of most of the parameters (range of overlap coefficient: [0.83,0.90]; Fig. 2). Pairwise comparisons of model parameters from the same origin show that most parameters did not shift in a consistent direction (Appendix S2 Tables 1). In contrast, *C. baileyi* parameter estimates tended to differ between models fit to data on control and removal plots, with parameter estimates for the environmental covariates showing relatively low overlap (range of overlap coefficient: [0.15-0.56]). Autoregressive terms, on the other hand, exhibited more overlap (AR (1) = 0.69, AR (12) = 0.67; Fig. 2). *C. baileyi* also exhibited high proportions of shifts in one direction for all environmental variables and the intercept (Appendix S2 Table 1). This suggests that the form of the forecasting model is dependent on the biotic context for this species.

**Figure 2.**
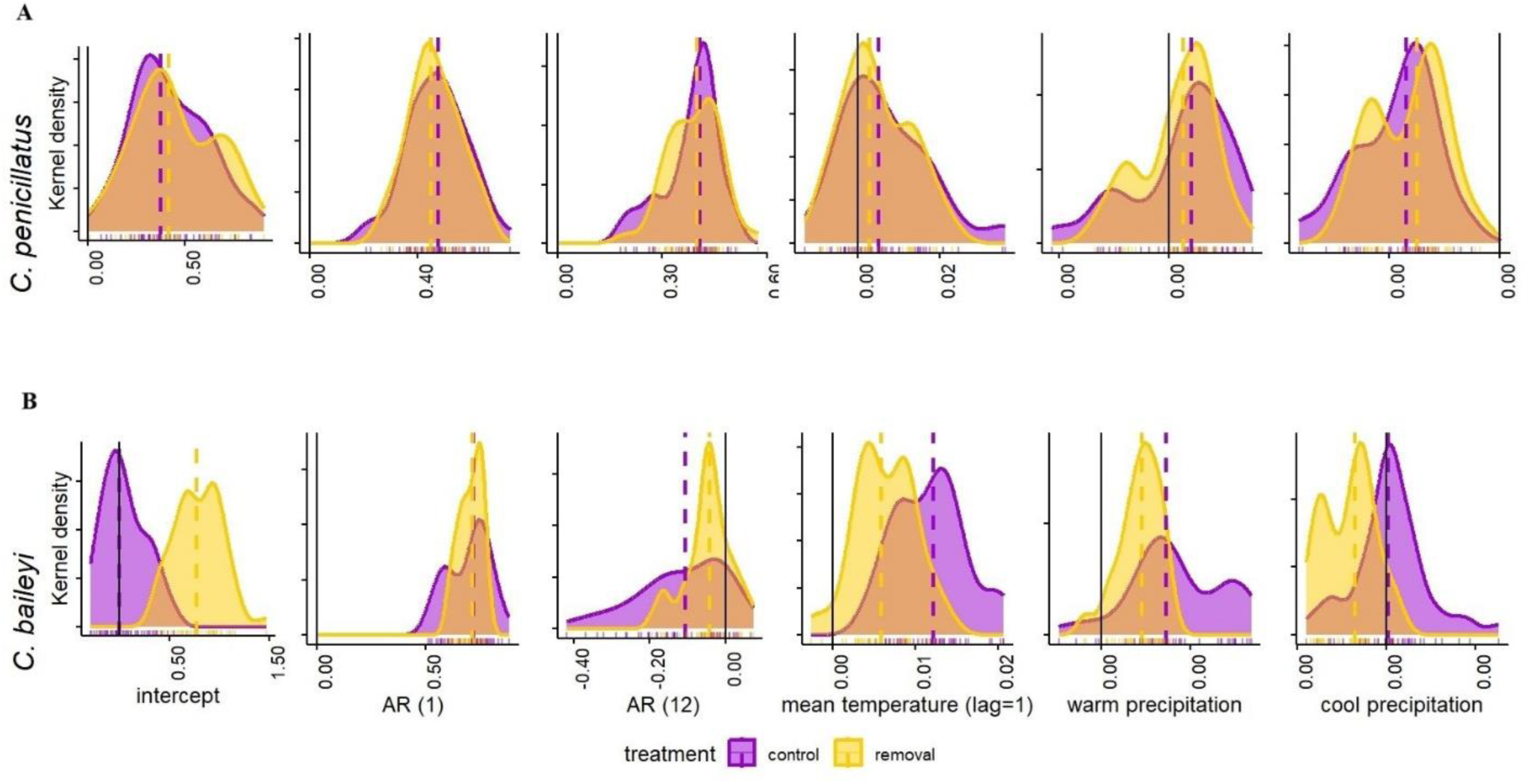
Kernel densities of the distribution of estimated coefficients (point estimates) obtained from the suite of rolling origin models fit to data on *C. penicillatus* (A; n = 45) and *C. baileyi* (B; n = 48) on control plots (purple densities) and removal plots (yellow densities).

### Model transferability under novel biotic conditions

The two species also differed in how well transferred models performed at forecasting compared to the non-transferred models. For *C. penicillatus* the transferred models performed similarly to the non-transferred models on both control and removal plot data (Figs. 3 and 4). Both transferred and non-transferred models showed a consistent pattern of increasing prediction error with increasing forecast horizon length for both RMSE and Brier score (Figs. 4A and 4B). In contrast, for *C. baileyi,* the transferred models generally performed worse than the non-transferred models when making forecasts (Fig. 3). Point forecast (RMSE) scores showed a clear pattern of better performance for the non-transferred models for both control and removal data (Figs. 4C and 4D). Brier scores were also generally better for the non-transferred model, particularly on the removal plots. However, the Brier score result was weaker when evaluating forecasts made for the control data. While the majority of origins showed worse forecasts for the transferred model, the mode of difference between the original and transferred model was near zero for all horizons (Fig. 4). This suggests higher uncertainty (as indicated by Brier score) in the less accurate predictions (as indicated by RMSE) from the models transferred from the removal plots to the control plots. Finally, similar to the *C. penicillatus* models, both models fit to *C. baileyi* data exhibited decreasing model performance at increasing horizons (Fig. 4).

**Figure 3.**
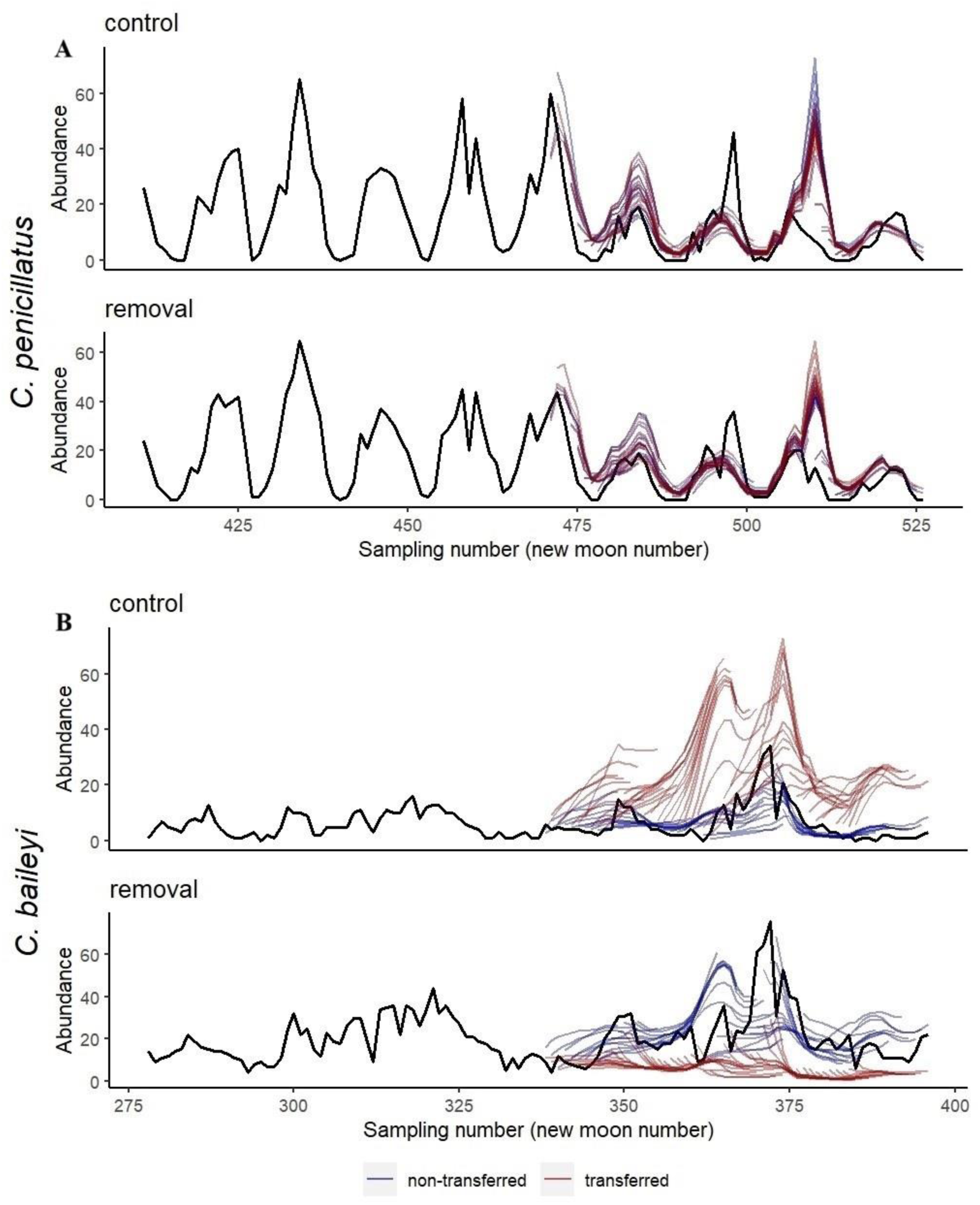
Predictions for *C. penicillatus* (A) and *C. baileyi* (B) abundances from models fit to non-transferred (blue lines) and transferred (red lines) data.

**Figure 4.**
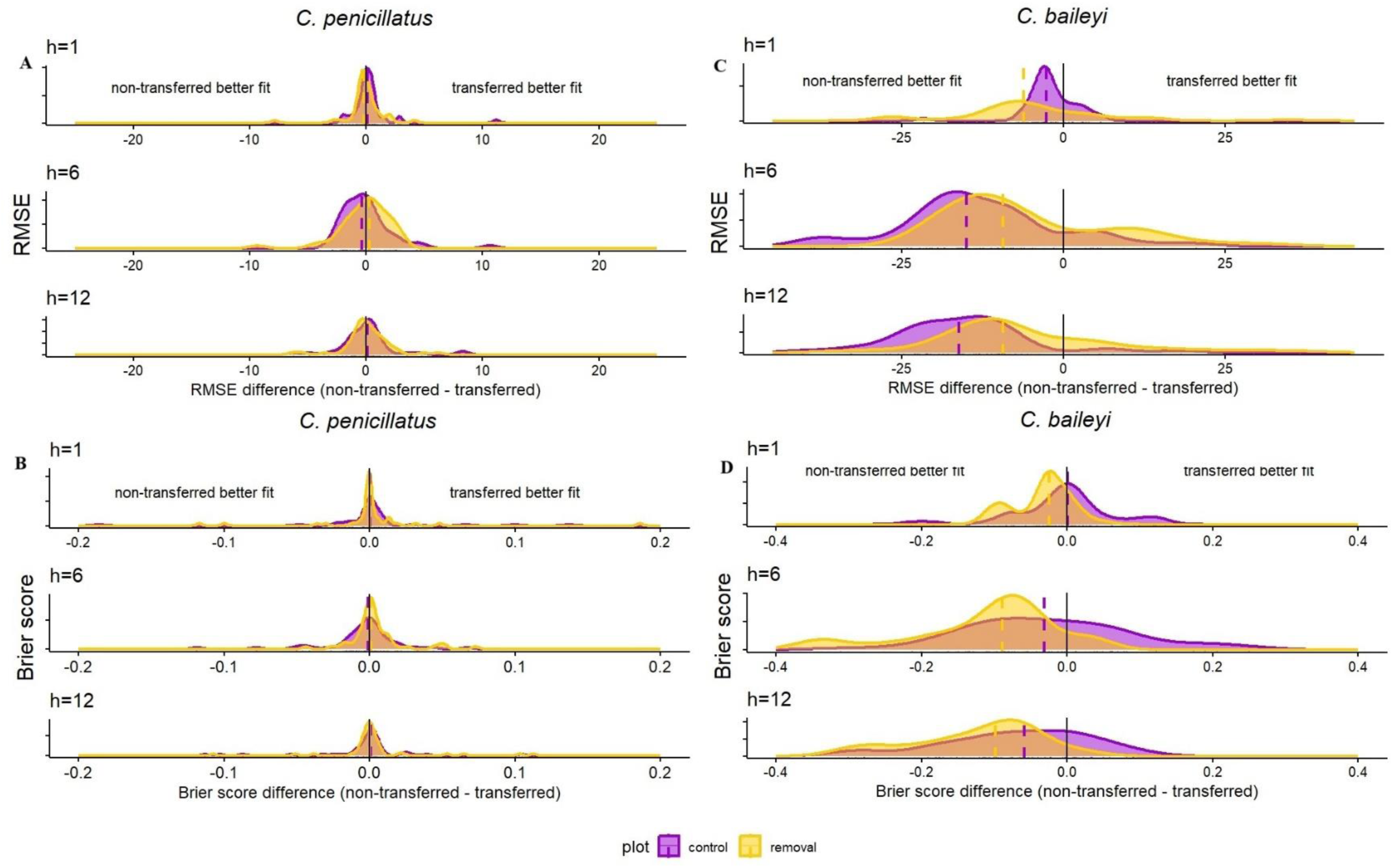
Difference of Root Mean Squared Error (RMSE) and Brier score of non-transferred and transferred models of *C. penicillatus* (A, B) and *C. baileyi* (C, D) on control (purple) and removal plots (yellow) at different forecast horizons.

## DISCUSSION

Ecological forecasts can be less accurate when models are transferred to novel biotic contexts. In this study, we observed this effect even though the long-term experimental nature of the Portal Project meant that plots with different species compositions were intermingled and there was no difference in the environmental conditions between the different biotic contexts. Decreased performance from models transferred to novel biotic conditions, however, depended on the species being forecast, with *C. baileyi* exhibiting significant shifts in both model parameters and forecast abundances, but *C. penicillatus* showing little change in either. This highlights the need to account for biotic interactions in forecasting models,and the need to understand when and why changes in the biotic context impact model transferability.

We expect changes in the biotic context to impact forecasting models if shifts in the biotic context involve species with strong influences on the species being forecast. *C. baileyi*, which colonized the site in 1995, increased in abundance when *Dipodomys* were experimentally removed, demonstrating strong competitive impacts by *Dipodomys* (Ernest and Brown 2001, Bledsoe and Ernest 2019). Our forecast models reflect this competitive impact with higher intercepts for the removal models when compared to the controls, which causes some of the divergence in forecasts when models are transferred. While *C. baileyi* is competitively inferior to *Dipodomys*, it is larger and competitively dominant over its congener *C. penicillatus*. With the removal of *Dipodomys*, *C. baileyi* shifts its stature in the community to that of the competitive dominant, which increases not only its abundance but could allow *C. baileyi* to exploit regions of its fundamental niche that overlap with *Dipodomys* (Thibault et al 2010, Diaz and Ernest 2022). A shift in the realized niche could explain differences in environmental parameters between control and removal plots as *C. baileyi* is no longer constrained by competition and can react more directly to the environmental drivers. The dual effect of altered competition on the intercept and environmental parameters then causes the transferred models to perform poorly (Fig. 2).

Changes in biotic context do not always alter competitive hierarchies, however. Like *C. baileyi*, *C. penicillatus* increases in abundance when *Dipodomys* spp. are removed, indicating a strong competitive interaction between these species (Valone and Brown 1995, Bledsoe and Ernest 2019, Diaz and Ernest 2022). With the establishment of *C. baileyi* on removal plots, however, the competitive pressures on *Dipodomys* removal plots increased. In response, *C. penicillatus* decreased their residency in the previously preferred plot (i.e., removals) and increased their probability of dispersing to nearby control plots (Bledsoe and Ernest 2019). Perhaps due to behavioral interactions between these territorial species, the less dominant *C. penicillatus* exhibited shifts in abundance between plots even when *C. baileyi* abundances decreased in 2010 (Bledsoe and Ernest 2019, Christensen et al 2019). Thus, *C. penicillatus* may perceive competition with its close congener as being a similar competitive environment to plots containing *Dipodomys* spp. This could explain the similarities in both the intercepts and the environmental parameters because competitive pressures are never alleviated and *C. penicillatus* has little opportunity to exploit unexpressed areas of its fundamental niche. Species with many weak interactions seem to be more forecastable (Durgaard et al 2022) as changes in a single competitor in the network are unlikely to result in a large shift in the expressed niche of the focal species being forecast. The fact that *C. penicillatus* does not exhibit significantly different dynamics despite the removal of *Dipodomys* highlights the challenges of understanding when biotic context will influence ecological forecasting due to complex species networks in nature.

Declines in the accuracy of forecasts with increasing forecast horizon exhibited an interesting interaction with model transfer to novel biotic contexts (Fig. 4). Decreasing forecast performance as forecasts are made further into the future is a common pattern in ecological forecasts (Dietze et al. 2018, Harris et al. 2018) that is demonstrated by both *C. baileyi* and *C. penicillatus* models. However, transferred models for *C. baileyi* decrease in forecast accuracy more rapidly with the forecast horizon (as indicated by increasing deviations between the original and transferred models, Figs. 4C and 4D). At short timescales, the strong short-term autoregressive components in the models allow good predictions even when transferring the model, but as the forecast horizon increases the differences in other model parameters become more prevalent leading to greater decay in accuracy for the transferred models (Fig. 4). This interaction suggests that estimates of decay in forecast accuracy may be overly optimistic if the composition of the community is also shifting at the timescales of the forecasts. This lends experimental support to the idea that estimates of model transferability need to consider multiple aspects of transfer (Gavish et al. 2017), in this case including both transfer to novel biotic context and transfer outside of the historical window used for fitting the models.

Differences between our two metrics for assessing forecast performance (RMSE and Brier score) demonstrate the importance of incorporating uncertainty in forecast evaluation and show an interesting interaction between uncertainty and model transfer to novel biotic contexts (Fig. 4). The RMSE, which only evaluates point estimates (not uncertainty), was worse for transferred *C. baileyi* models on both control and removal plots, even at short horizons. The Brier score, which integrates model uncertainty, exhibited a similar pattern for the removal plot data, but showed reduced responses to model transfer on the control plots (Fig. 4D). This difference between the Brier score and RMSE response suggests that while the predictions from the transferred removal models are less accurate, the uncertainty in those predictions is also higher, so the model is less confident in the less accurate predictions. Potentially, models fit to the removals exhibit better uncertainty under model transfer because these models are exposed to a wider range of variation in abundance than models fit to the control plots. Due to the competitive release from *Dipodomys spp.*, *C. baileyi* abundances are typically higher and more variable in removal plots. This wider range of variation is likely due to reduced constraints on population growth during good years and potentially a shift in response to environmental drivers. If this increased variation is not fully captured by the models’ dynamics it will result in increased error terms and uncertainty, thus resulting in predictions that are penalized less by evaluation metrics that include uncertainty. This complex interaction between model transfer, uncertainty, and experimental treatment suggests that it is important to incorporate uncertainty into the assessment of model transferability because it can provide insights that are different from point estimates alone. It also shows that, in some cases, transferred models may be appropriate for decision making even if they make less accurate point forecasts, as long as the decision making properly incorporates uncertainty. In general, evaluating uncertainty - either by using metrics that include it or by measuring model transferability and associated forecast uncertainties - will be important for assessing how effectively models can be transferred and their utility for implementing conservation strategies on species or locations with limited data availability (Houlahan et al. 2017, Yates et al. 2018).

In this study, we focused on single species models to demonstrate and assess model transferability under varying biotic conditions. Single species models are common in ecological modeling, forecasting, and management, but because they do not attempt to model species interactions these models are likely to be particularly susceptible to changes in the biotic context. Multivariate community models, which can include species interactions, have the potential to provide improved transfer to novel biotic conditions by incorporating information on processes such as competition. For example, for the control plots this type of model could include the interactions between *C. baileyi* and *Dipodomys* species, potentially allowing it to transfer more effectively to the removal plots where *Dipodomys* abundance would influence predictions as an observed value at or near zero. The use of these types of models in dynamic ecological forecasting remains uncommon since the number of ecosystems with sufficiently long time-series on all of the key species in the community is limited. Since explicitly modeling interactions is important for modeling population dynamics (e.g., Lima et al. 2008), species distributions (e.g., Pollock et al. 2014), and model transferability, further exploration of multivariate community predictions will be an important next step for ecological forecasting.

We have shown that changes in the presence of other species can impact both the parameters of ecological forecasting models and their predictions. This suggests that caution will be necessary when making forecasts in new systems or over long enough periods of time that the composition of other species in the community undergoes change. This is important because the development of ecological forecasting models for new scenarios is often limited by data availability. Transferring models to new scenarios, using either no new data or a small amount of new data to fine-tune the model, is a common approach to addressing this challenge (Houlahan et al. 2017, Yates et al. 2018, Lewis et al. 2023). Therefore, models that better represent the complex dynamics of biological interactions, and effectively predict beyond the conditions they were built on, are needed in an era of fast-paced environmental change (Yates et al. 2018). Developing models that can be effectively transferred across space, time, and biotic context, and effectively communicate the uncertainties in their predictions, are important endeavors to facilitate the expanded development and use of ecological forecasts (Houlahan et al. 2017, Yates et al. 2018).

## ACKNOWLEDGMENTS

We thank the volunteers, staff, and research assistants who collected the Portal Project data over the years. This work was supported by research grants awarded to SKM Ernest and EP White by the US National Science Foundation (NSF DEB-1929730), USDA National Institute of Food and Agriculture, Hatch Project FLA-WEC-005983 (Ernest) and FLA-WEC-005944 (White).

