## Appendix S1 for "Transferability of ecological forecasting models to novel biotic conditions in a long-term experimental study"

**Journal**: Ecology

**Manuscript type**: Article

Ernest, and Ethan P. White

### **APPENDIX S1**

#### **METHODS**

##### **Collinearity among predictors and estimated parameters**

We performed pairwise correlation tests on all possible pairs of the environmental covariates used in our models (i.e., mean temperature (lag=1), warm and cool precipitation) using Pearson’s correlation test. Then, we assessed the collinearity of the estimated parameters from each treatment-specific model by conducting covariance and correlation assessments on the estimated parameters generated from sequential model fitting. For each model fit at each origin, we computed a covariance matrix from a given Fisher information matrix by inversion using the invertinfo() function in the ‘tscount’ package. Collinearity was low for the covariates and their estimated parameters (Appendix S1 Figs.1-5).

#### **TABLES**

Table 1. Akaike Information Criterion (AIC) scores of Negative Binomial and Poisson models fit to data on desert and Bailey’s pocket mice on control and removal plots. The use of the negative binomial model is strongly supported by the difference in AIC values.

| Species | Control | | Removal | |
| --- | --- | --- | --- | --- |
|  | Negative Binomial | Poisson | Negative Binomial | Poisson |
| Desert pocket mouse | 764.64 | 975.39 | 755.09 | 895.17 |
| Bailey’s pocket mouse | 577.15 | 600.60 | 800.01 | 874.72 |

### **APPENDIX S1 FIGURES**

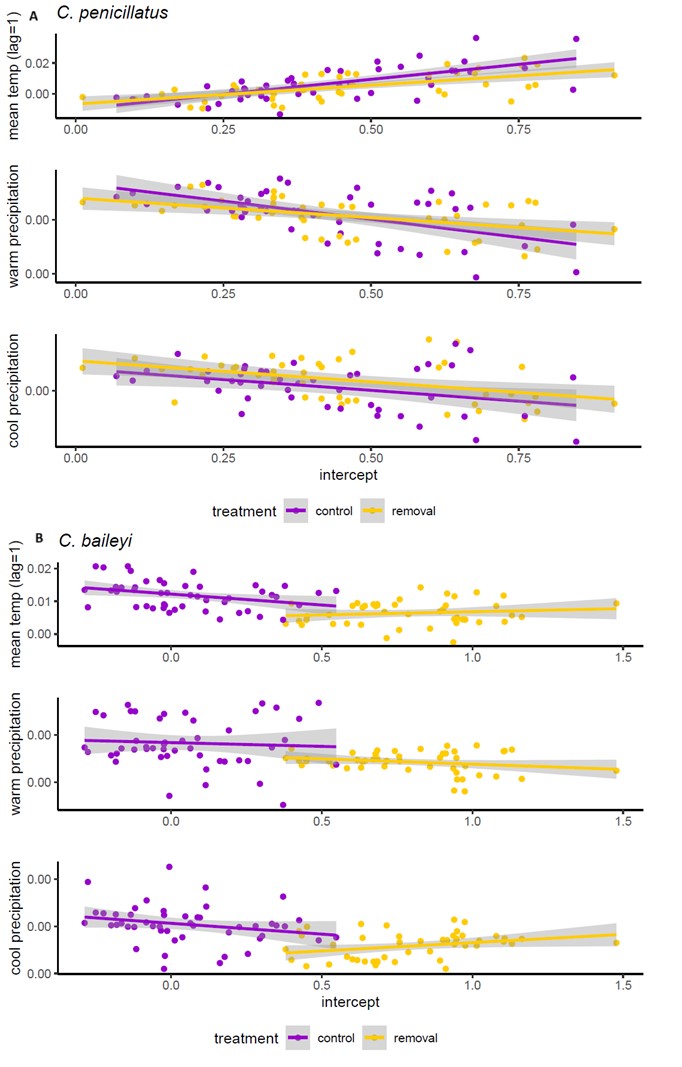

**Appendix S1 Fig. 1.** Covariances of the intercept and the slopes at different origins of time-series models fit to data on *Chaetodipus penicillatus* (A) and *C. baileyi* (B) in control and removal plots.

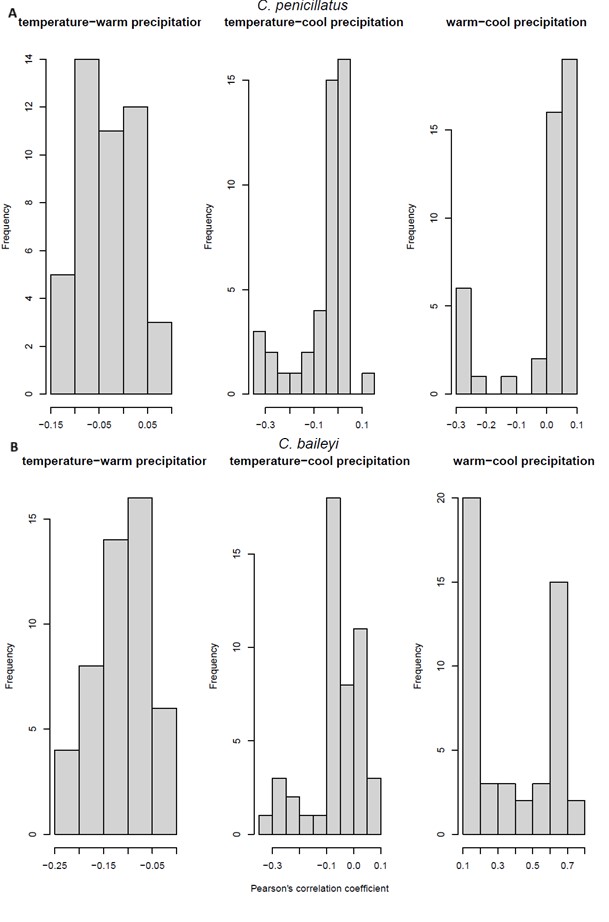

**Appendix S1 Fig. 2.** Frequency of correlation coefficients of environmental covariates. Correlation coefficients on the raw values of the environmental covariates used in models fit to data on *Chaetodipus* *penicillatus* (A) and *C. baileyi* (B) in a long-term experiment in Portal, AZ**.**

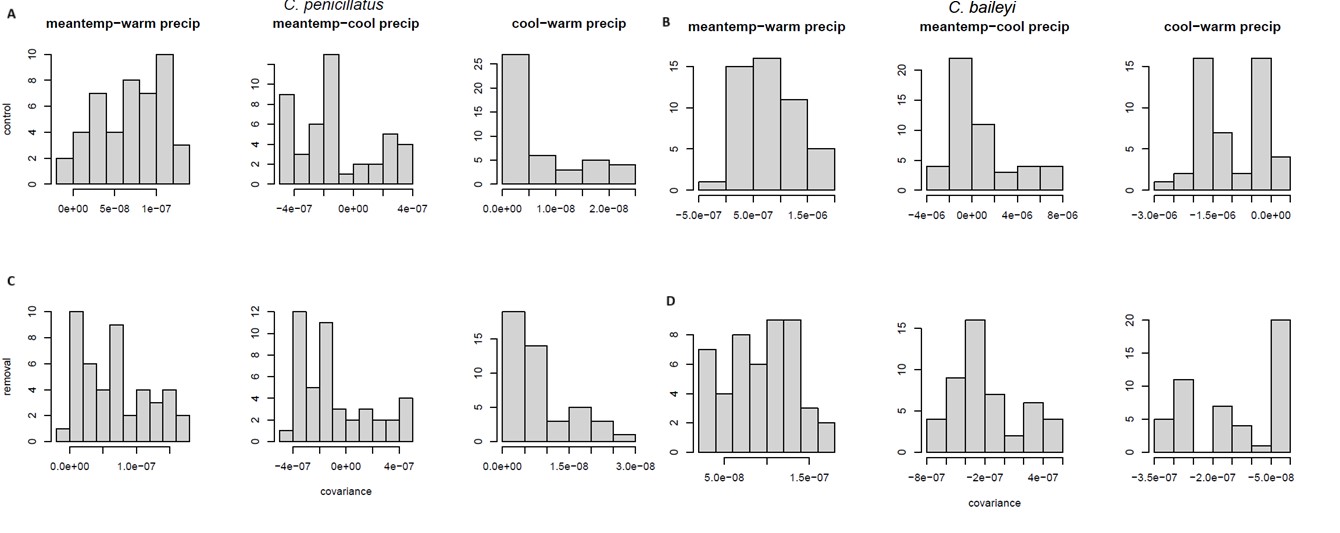

**Appendix S1 Fig. 3.** Pairwise covariance of environmental parameter estimates. Estimates obtained from time-series models on *C. penicillatus* in control (A) and removal plots (C) and on *C. baileyi* in control (B) and removal (D) plots.

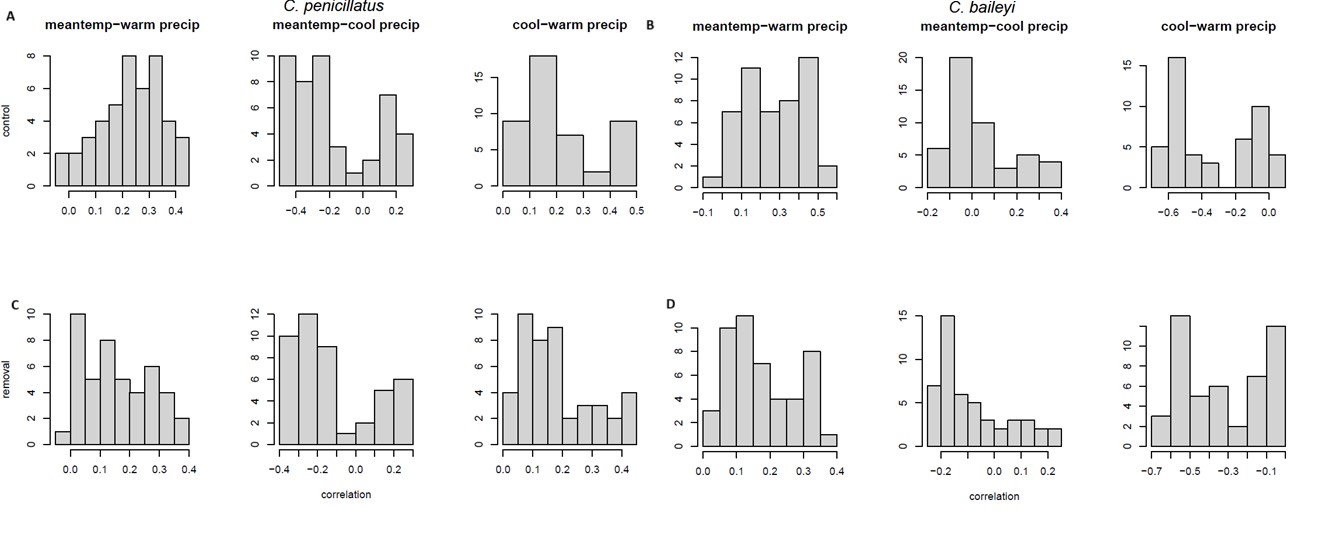

**Appendix S1 Fig. 4.** Pairwise correlation of environmental parameter estimates. Correlation estimates obtained from time-series models on *C. penicillatus* in control (A) and removal (C) plots and *C. baileyi* in control (B) and removal (D) plots.

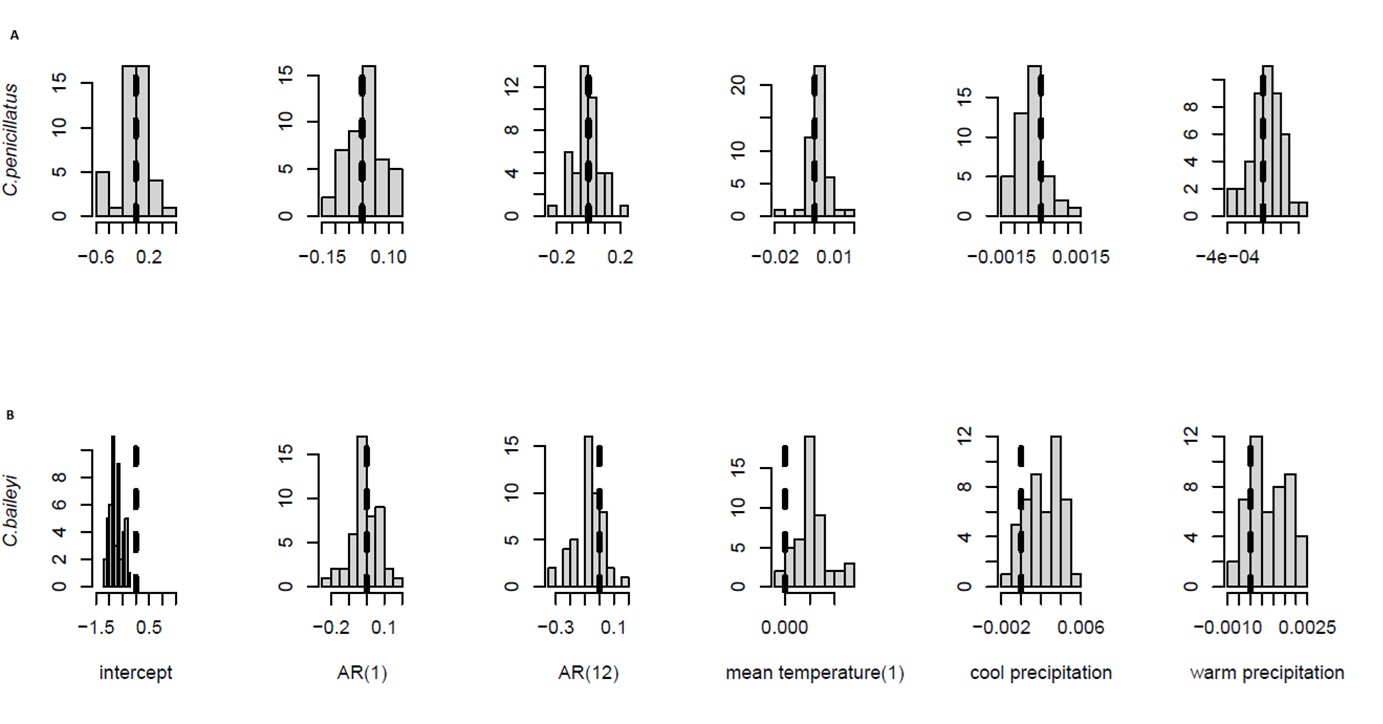

**Appendix S1 Fig. 5.** Distribution of pairwise changes in parameter estimates (difference in parameter estimates generated from control and removal models) fit to *C. penicillatus* (A) and *C. baileyi* (B) data.

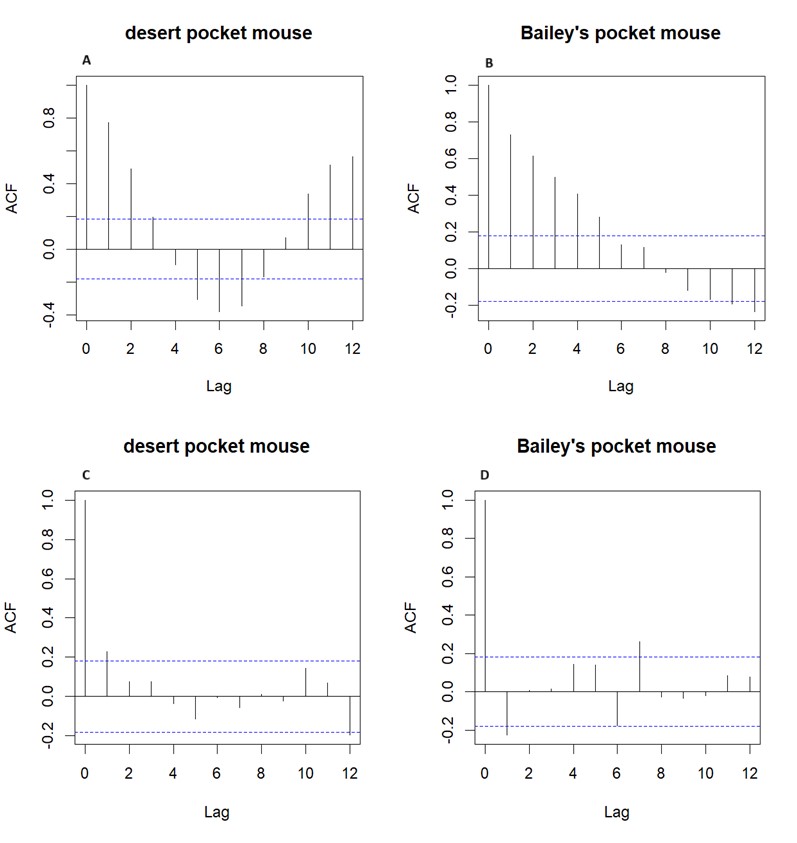

**Appendix S1 Fig. 6**. Autocorrelation function (ACF) of count data on Bailey’s and desert pocket mouse. ACF on the control plot data for desert (A) and Bailey’s (B) pocket mice, and ACF on model residuals for models fit to the full control time-series for the desert (C) and the Bailey’s pocket mouse (D).
