## Appendix S2 for "Transferability of ecological forecasting models to novel biotic conditions in a long-term experimental study"

**Journal**: Ecology

**Manuscript type**: Article

### APPENDIX S1

#### METHODS

***Collinearity among predictors and estimated parameters***

We performed pairwise correlation tests on all possible pairs of the environmental covariates used in our models (i.e., mean temperature (lag=1), warm and cool precipitation) using Pearson's correlation test. Then, we assessed the collinearity of the estimated parameters from each treatment-specific model by conducting covariance and correlation assessments on the estimated parameters generated from sequential model fitting. For each model fit at each origin, we computed a covariance matrix from a given Fisher information matrix by inversion using the invertinfo() function in the 'tscount' package. Collinearity was low for the covariates and their estimated parameters (Appendix S1 Figs.1-5).

**TABLES**

Table 1. Akaike Information Criterion (AIC) scores of Negative Binomial and Poisson models fit to data on desert and Bailey's pocket mice on control and removal plots. The use of the negative binomial model is strongly supported by the difference in AIC values.

| Species | Control | | Removal | |
|---|---|---|---|---|
| | Negative Binomial | Poisson | Negative Binomial | Poisson |
| Desert pocket mouse | 764.64 | 975.39 | 755.09 | 895.17 |
| Bailey's pocket mouse | 577.15 | 600.60 | 800.01 | 874.72 |

### APPENDIX S1 FIGURES

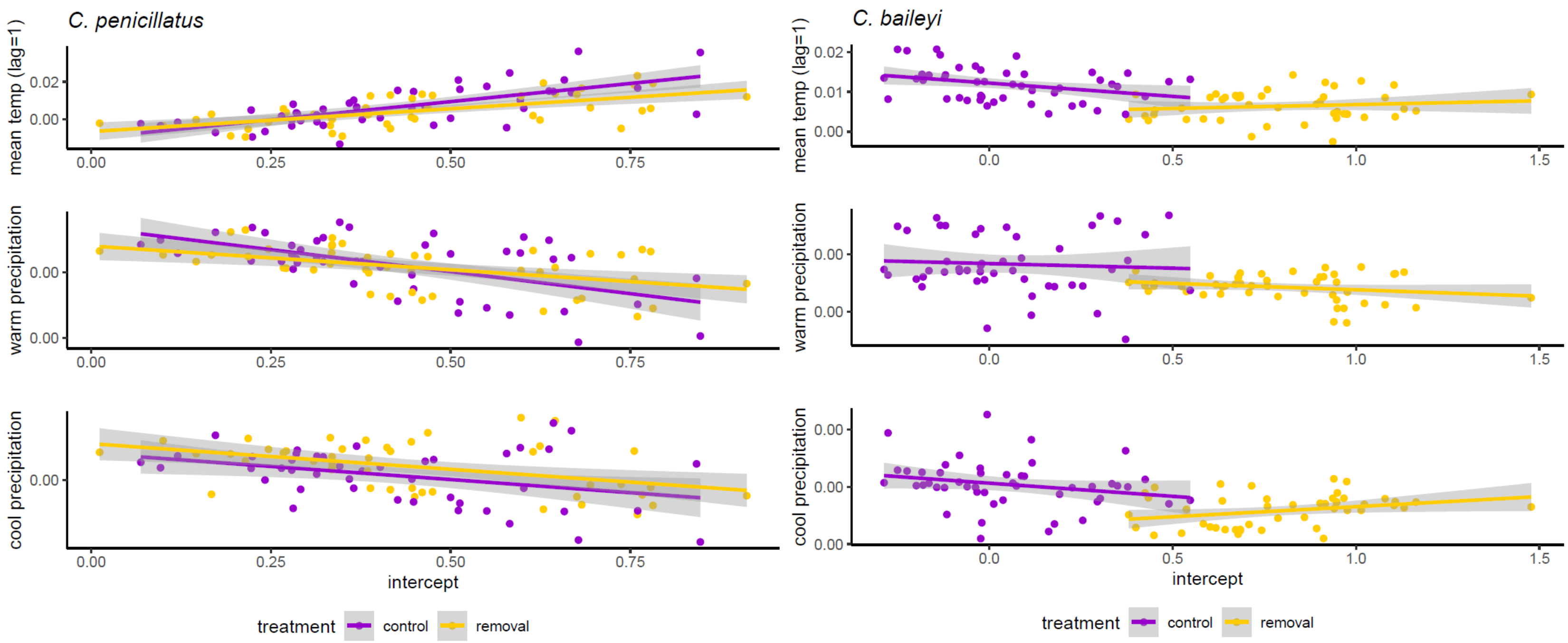


**Appendix S1 Fig. 1.** Covariances of the intercept and the slopes at different origins of time-series models fit to data on *Chaetodipus penicillatus* (left panel) and *C. baileyi* in control and removal plots.

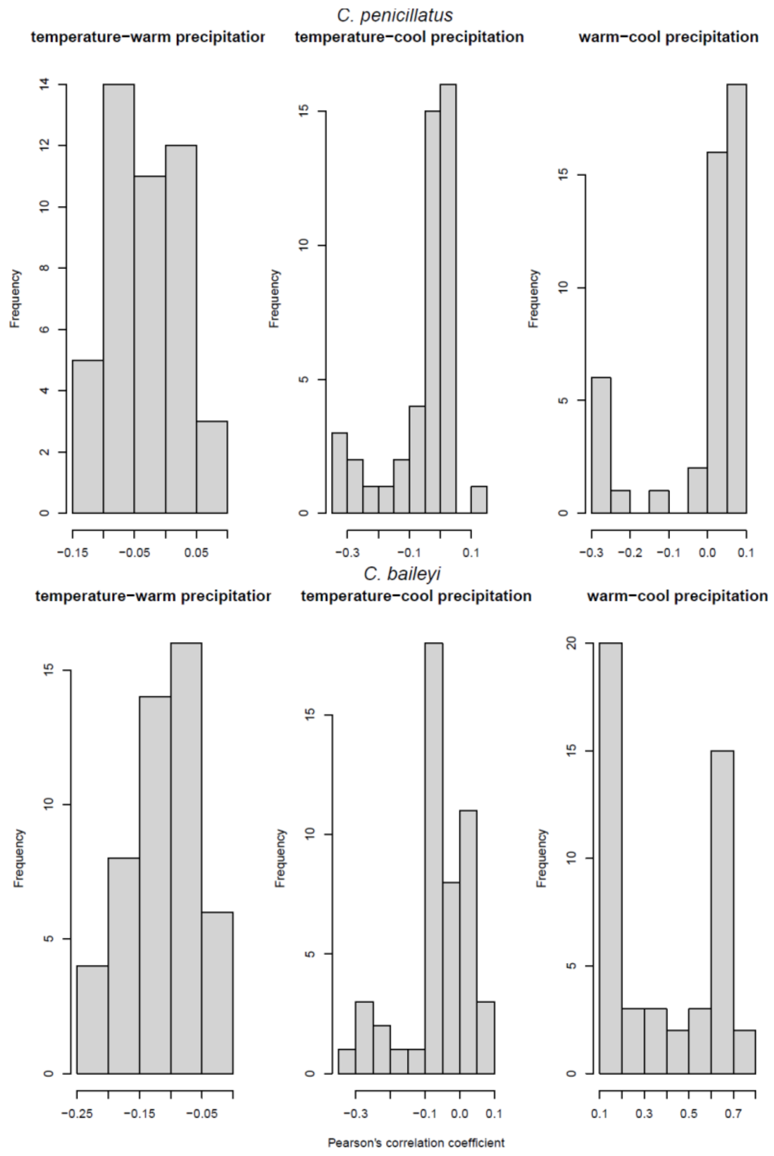


**Appendix S1 Fig. 2.** Frequency of the correlation coefficients on the raw values of the environmental covariates used in models fit to data on *Chaetodipus penicillatus* (left panel) and *C. baileyi* (right panel) in a long-term experiment in Portal, AZ.

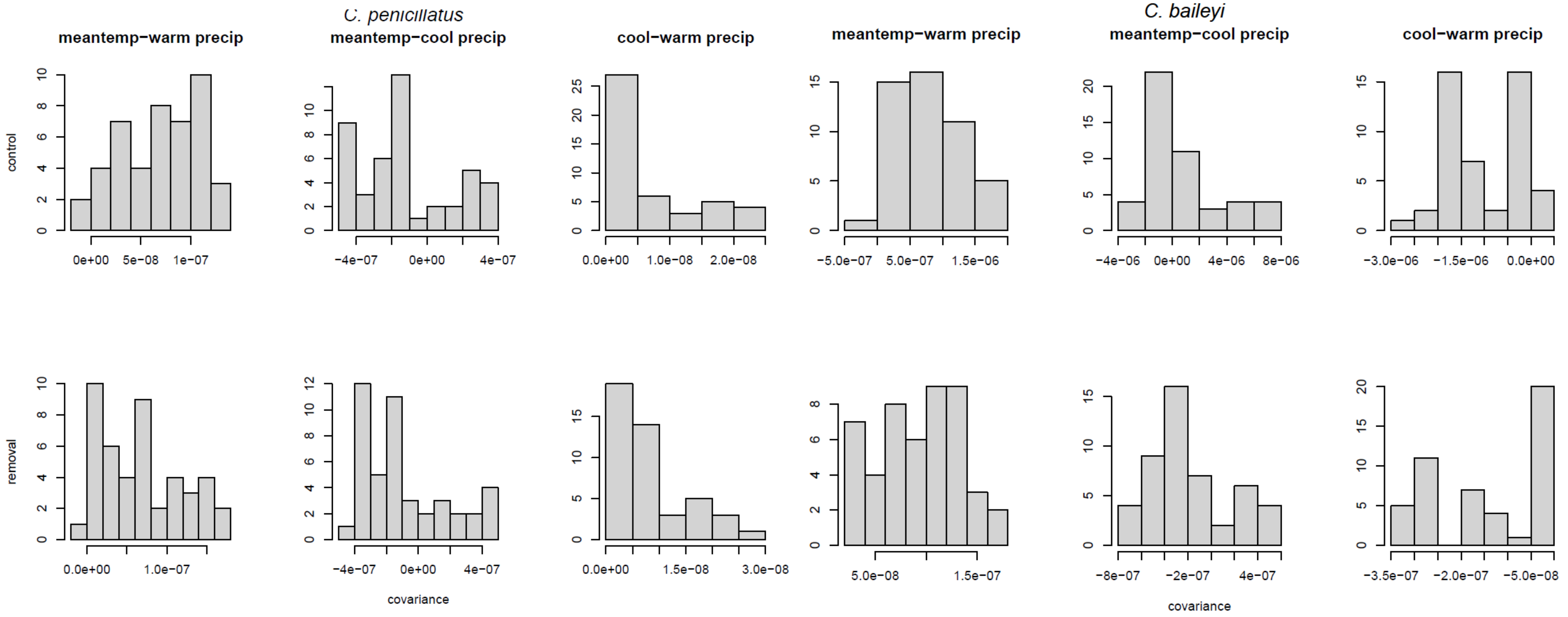


**Appendix S1 Fig. 3.** Pairwise covariance of the environmental parameter estimates obtained from time-series models on *C. penicillatus* (left panel) and *C. baileyi* (right panel) on control (top panel) and removal (bottom panel) plots.

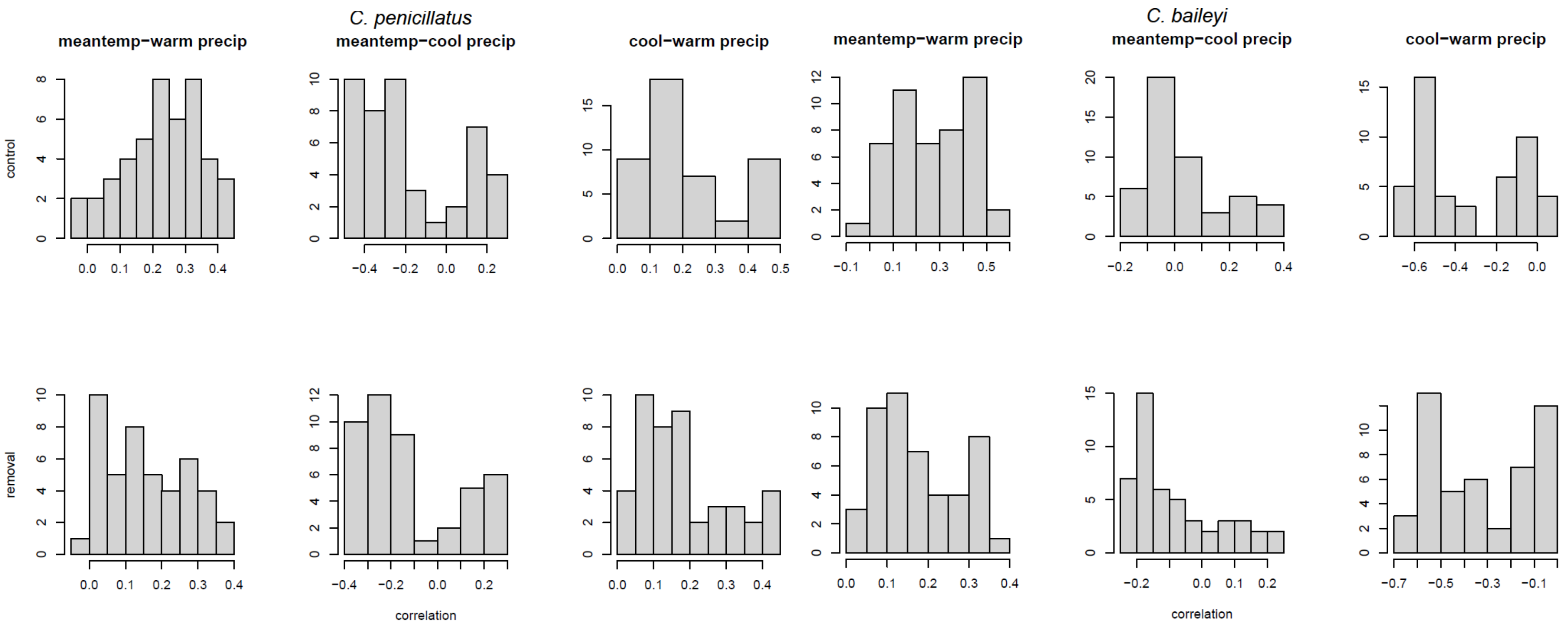


**Appendix S1 Fig. 4.** Pairwise correlation of the environmental parameter estimates obtained from time-series models on *C. penicillatus* (top panel; plots 1-3) and *C. baileyi* (bottom panel; plots 4-6) in control (left panel; plots 7-9) and removal (right panel; plots 10-12).

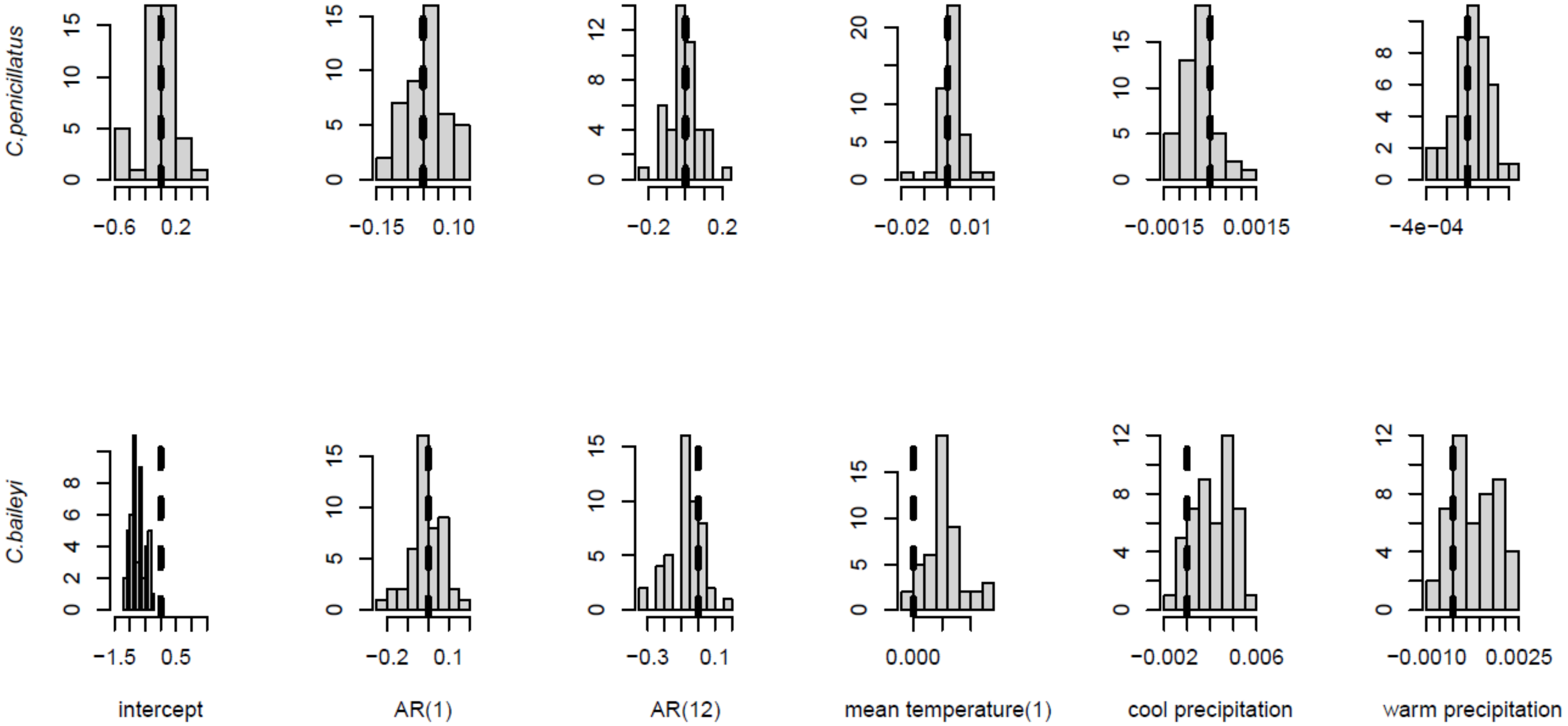


**Appendix S1 Fig. 5.** Distribution of pairwise changes in parameter estimates (difference in parameter estimates generated from control and removal models) fit to *C. penicillatus* (top panel) and *C. baileyi* (bottom panel) data.

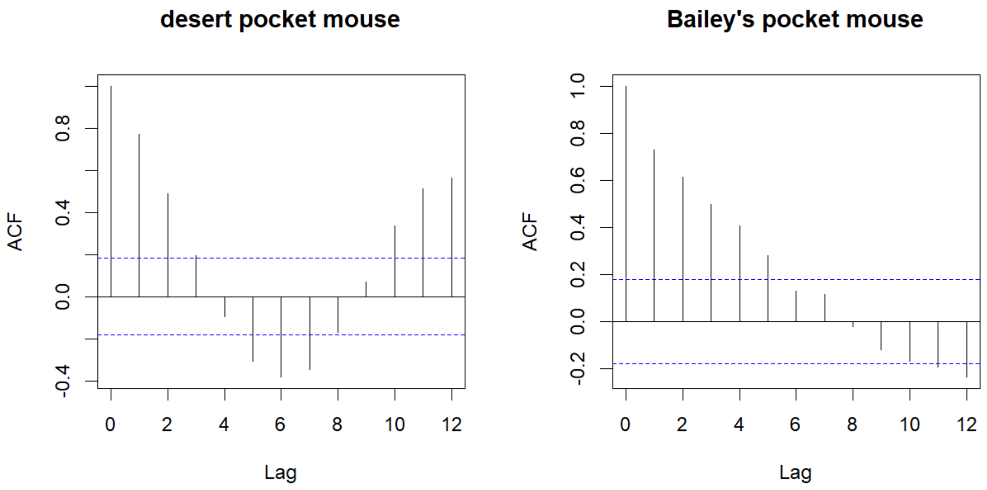


**Appendix S1 Fig. 6**. Autocorrelation function (ACF) of count data on the control plots for the desert pocket mouse (top panel) and the Bailey's pocket mouse (bottom panel).

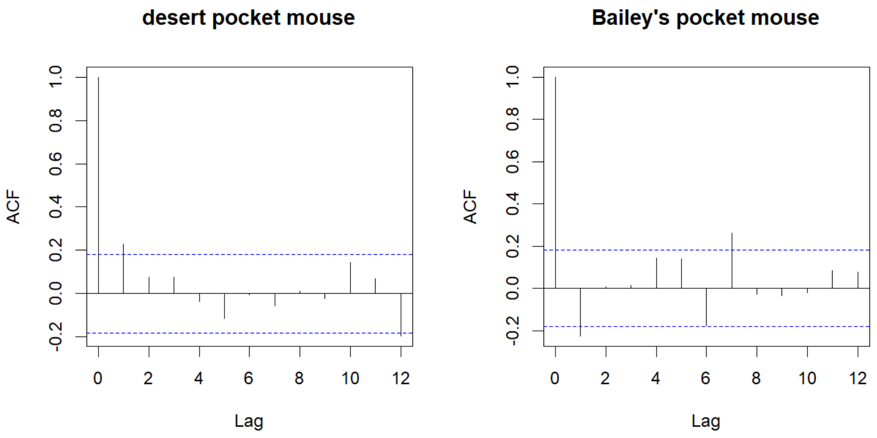


**Appendix S1 Fig. 7**. Autocorrelation function (ACF) of the model residuals for models fit to the full control time-series for the desert pocket mouse (left) and Bailey's pocket mouse (right).
